# HiCRep: assessing the reproducibility of Hi-C data using a stratum-adjusted correlation coefficient

**DOI:** 10.1101/101386

**Authors:** Tao Yang, Feipeng Zhang, Galip Gürkan Yardımcı, Fan Song, Ross C. Hardison, William Stafford Noble, Feng Yue, Qunhua Li

## Abstract

Hi-C is a powerful technology for studying genome-wide chromatin interactions. However, current methods for assessing Hi-C data reproducibility can produce misleading results because they ignore spatial features in Hi-C data, such as domain structure and distance dependence. We present HiCRep, a framework for assessing the reproducibility of Hi-C data that systematically accounts for these features. In particular, we introduce a novel similarity measure, the stratum adjusted correlation coefficient (SCC), for quantifying the similarity between Hi-C interaction matrices. Not only does it provide a statistically sound and reliable evaluation of reproducibility, SCC can also be used to quantify differences between Hi-C contact matrices and to determine the optimal sequencing depth for a desired resolution. The measure consistently shows higher accuracy than existing approaches in distinguishing subtle differences in reproducibility and depicting interrelationships of cell lineages. The proposed measure is straightforward to interpret and easy to compute, making it well-suited for providing standardized, interpretable, automatable, and scalable quality control. The freely available R package HiCRep implements our approach.

## Introduction

The three-dimensional (3D) genome organization across a wide range of length scales is important for proper cellular functions (Dekker et al. 2013; Sexton and Cavalli 2015; Bickmore 2013). At large distances, non-random hierarchical territories of chromosomes inside the cell nucleus are tightly linked with gene regulation (Misteli 2010). At a finer resolution, the interactions between distal regulatory elements and their target genes are essential for orchestrating correct gene expression across time and space (e.g. different tissues). A progression of high-throughput methods based on chromatin conformation capture (3C) (Dekker 2002) has emerged, including 4C (Simonis et al. 2006), 5C (Dostie et al. 2006), Hi-C (Lieberman-aiden et al. 2009), ChIA-PET (Li et al. 2010), Capture Hi-C (Hughes et al. 2014; Mifsud et al. 2015), and HiChIP (Mumbach et al. 2016). These methods offer an unprecedented opportunity to study higher-order chromatin structure at various scales. Among them, the Hi-C technology and its variants are of particular interest due to their relatively unbiased genome-wide coverage and ability to measure chromatin interaction intensities between any two given genomic loci.

However, the analysis and interpretation of Hi-C data are still in their early stages. In particular, no sound statistical metric to evaluate the quality of Hi-C data has been developed. When biological replicates are not available, investigators often rely on either visual inspection of the Hi-C interaction heatmap or examination of the ratio of long-range interaction read pairs over the total sequenced reads (Dixon et al. 2012, 2015; Jin et al. 2013), but neither of these approaches is supported by robust statistics. When two or more biological replicates are available, it is a common practice to use either Pearson or Spearman correlation coefficients between the two Hi-C data matrices as a metric for data quality (Imakaev et al. 2012; Hu et al. 2012; Gorkin et al. 2014; Rao et al. 2014; Ay and Noble 2015; Servant et al. 2015; Dixon et al. 2015). However, Hi-C data have certain unique characteristics, including domain structures, such as topological association domain (TAD) and A/B compartments, and distance dependence, which refers to the fact that the chromatin interaction frequencies between two genomic loci, on average, decrease substantially as their genomic distance increases. Standard correlation approaches do not take into consideration these structures and may lead to incorrect conclusions. As we will demonstrate, two unrelated biological samples can have a high Pearson correlation coefficient, while two visually similar replicates can have a low Spearman correlation coefficient. It is also not uncommon to observe higher Pearson and Spearman correlations between unrelated samples than those between real biological replicates.

In this work, we develop HiCRep, a novel framework for assessing the reproducibility of Hi-C data that takes into account the unique spatial features of the data. HiCRep first minimizes the effect of noise and biases by smoothing the Hi-C matrix, and then it addresses the distance-dependence effect by stratifying Hi-C data according to their genomic distance. In particular, we develop a stratum-adjusted correlation coefficient (SCC) as a similarity measure of Hi-C interaction matrices. SCC shares the similar range and interpretation as the standard correlation coefficients, making it easily interpretable. It can be used to assess the reproducibility of replicate samples or quantify the distance between Hi-C matrices from different cell types. Our framework also estimates confidence intervals for SCC, making it possible to infer statistical significance of the difference in reproducibility measurements. We applied our method to three different groups of publicly available Hi-C data sets to illustrate its power in distinguishing subtle differences between closely related cell lines and biological replicates and resolving interrelationship between different cell lineages or tissues.

## Results

### Spatial patterns in Hi-C data and their influence on reproducibility assessment

Unlike many other genomic data types, Hi-C data exhibits unique spatial patterns. One prominent pattern is the strong decay of interaction frequency as genomic distance increases between interaction loci, i.e. the distance dependence. This pattern is generally thought to result from non-specific interactions, which are more likely to occur between loci at closer genomic distance than those at a greater distance (Lajoie et al. 2015; Fudenberg and Mirny 2012). It is found consistently in every Hi-C matrix and is one of the most dominant patterns in the matrix of interaction frequencies measured by Hi-C (Lajoie et al. 2015). This dependence on distance generates strong but spurious association between Hi-C matrices even when the samples are unrelated, as revealed by the high Pearson correlation between any two Hi-C matrices. As an example, we computed the Pearson correlations of Hi-C contact matrices between two biological replicates and between two unrelated cell lines, hESC and IMR90 (Dixon et al. 2012). Though samples from the same cell line are expected to be much more correlated to each other than samples from unrelated cell lines, the Pearson correlation shows little difference between samples from different cell types (a hESC sample and an IMR90 sample, ρ=0.92) and biological replicates in hESC (ρ = 0.91)(Fig. 1A). Further investigation shows that the dependence pattern between the contact intensity and distance (Fig. 1B) is highly similar in hESC and IMR90, which creates the high, spurious correlation between the Hi-C samples from these two cell lines. Therefore, the Pearson correlation coefficient cannot distinguish real biological replicates from unrelated samples.

**Figure 1.**
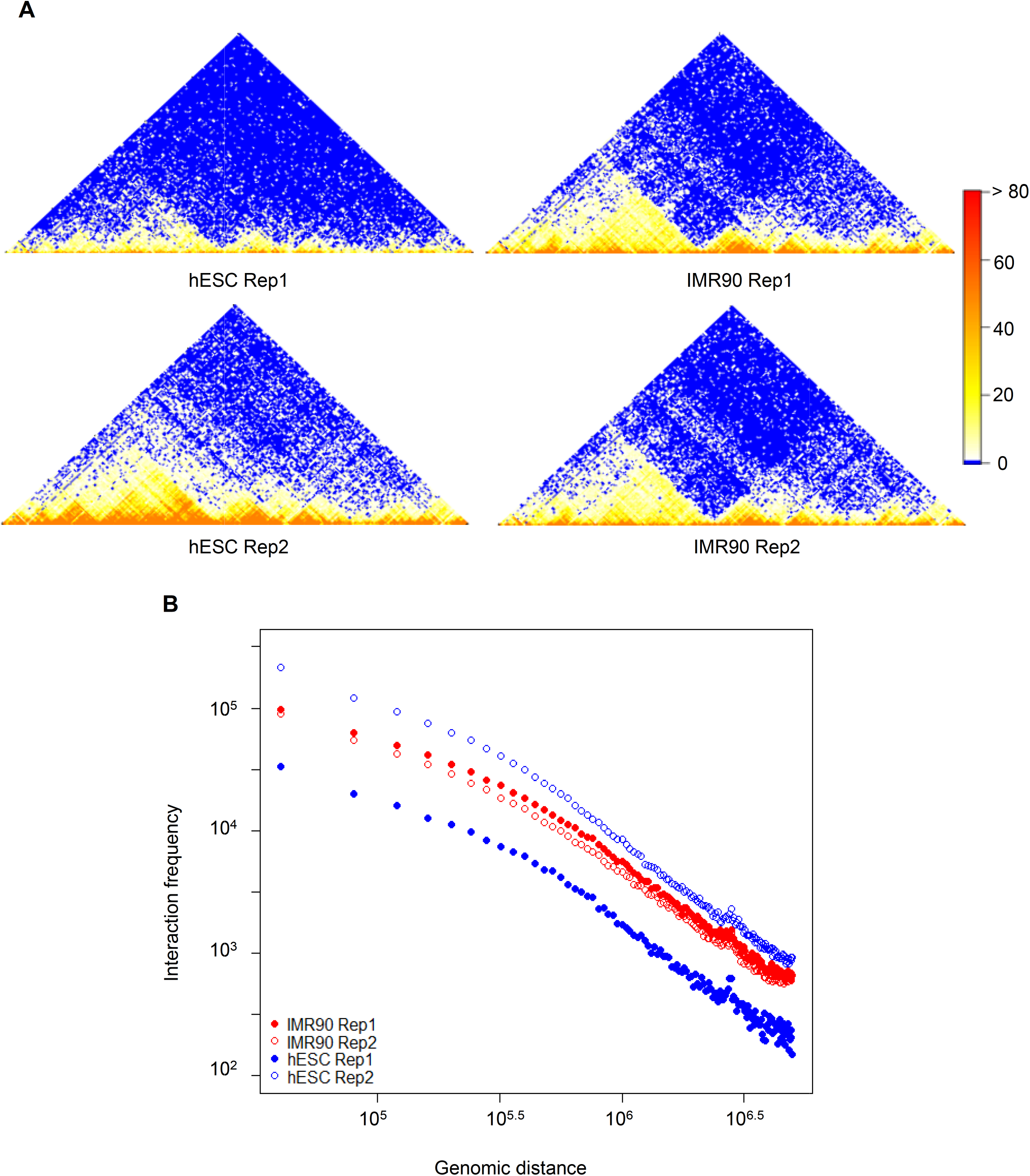
An illustration example. (A) Hi-C contact maps of the biological replicates of hESC and IMR90. (B) Relationship between genomic distance and the average contact frequency for the samples in (A). Data is from chromosome 22: 32000000 – 40000000.

Another important pattern of Hi-C data is the domain structure in contact maps. These structures represent contiguous regions in which loci tend to interact more frequently with each other than with outside regions. While the interactions within the structures can be highly variable between different cell types, the domain structures, such as topologically associating domains (TADs), are stable across cell types (Dixon et al. 2012; Rapkin et al. 2012; Nagano et al. 2013). Therefore, we expect a higher reproducibility at the domain level than at the individual contact level. This difference should be reflected in the reproducibility assessment. However, both Pearson and Spearman correlation coefficients only consider point interactions, and do not take domain structures into account. A consequence of this is that Spearman correlation can be driven to low values by the stochastic variation in the point interactions and overlook the similarity in domain structures. As a result, two biological replicates that have highly similar domain structures may have a low Spearman correlation coefficient; conversely, a sample may have a higher Spearman correlation with an unrelated sample than with its biological replicates when the stochastic variation is high. For instance, despite the high similarity between the biological replicates in IMR90 and hESC, their Spearman correlations are only 0.47 and 0.37, respectively. However, the Spearman correlation between an IMR90 sample and a hESC sample (0.44) is higher than the correlation between the two hESC replicates (0.37), even though there are many differences in the domain structures of the two cell lines. Therefore, we need a more sophisticated evaluation metric to incorporate both structural aspects of variation for a better assessment of the reproducibility of Hi-C data.

### Overview of the HiCRep method

We develop a novel two-stage approach to evaluate the reproducibility of Hi-C data (Fig. 2). The first stage is smoothing the raw contact matrix in order to reduce local noise in the contact map and to make domain structures more visible. The smoothing is accomplished by applying a 2D mean filter, which replaces the read count of each contact in the contact map with the average counts of all contacts in its neighborhood. In the second stage, we apply a stratification approach to account for the pronounced distance dependence in the Hi-C data. This stage proceeds in two steps. First we stratify the smoothed chromatin interactions according to their genomic distance, and then we apply a novel stratum-adjusted correlation coefficient statistic (SCC) to assess the reproducibility of the Hi-C matrices. The SCC statistic is calculated by computing a Pearson correlation coefficient for each stratum (Fig. 2) and then aggregating the stratum-specific correlation coefficients using a weighted average, with the weights derived from the generalized Cochran-Mantel-Haenszel (CMH) statistic (Mantel 1963; Agresti 2012). The value of SCC ranges from -1 to 1 and can be interpreted in a way similar to the standard correlation. A great advantage of our approach is that we can derive the asymptotic variance of SCC and use it to assess statistical significance when comparing reproducibility from different samples. More detailed descriptions of the HiCRep method and the SCC statistic are presented in the Methods section.

**Figure 2.**
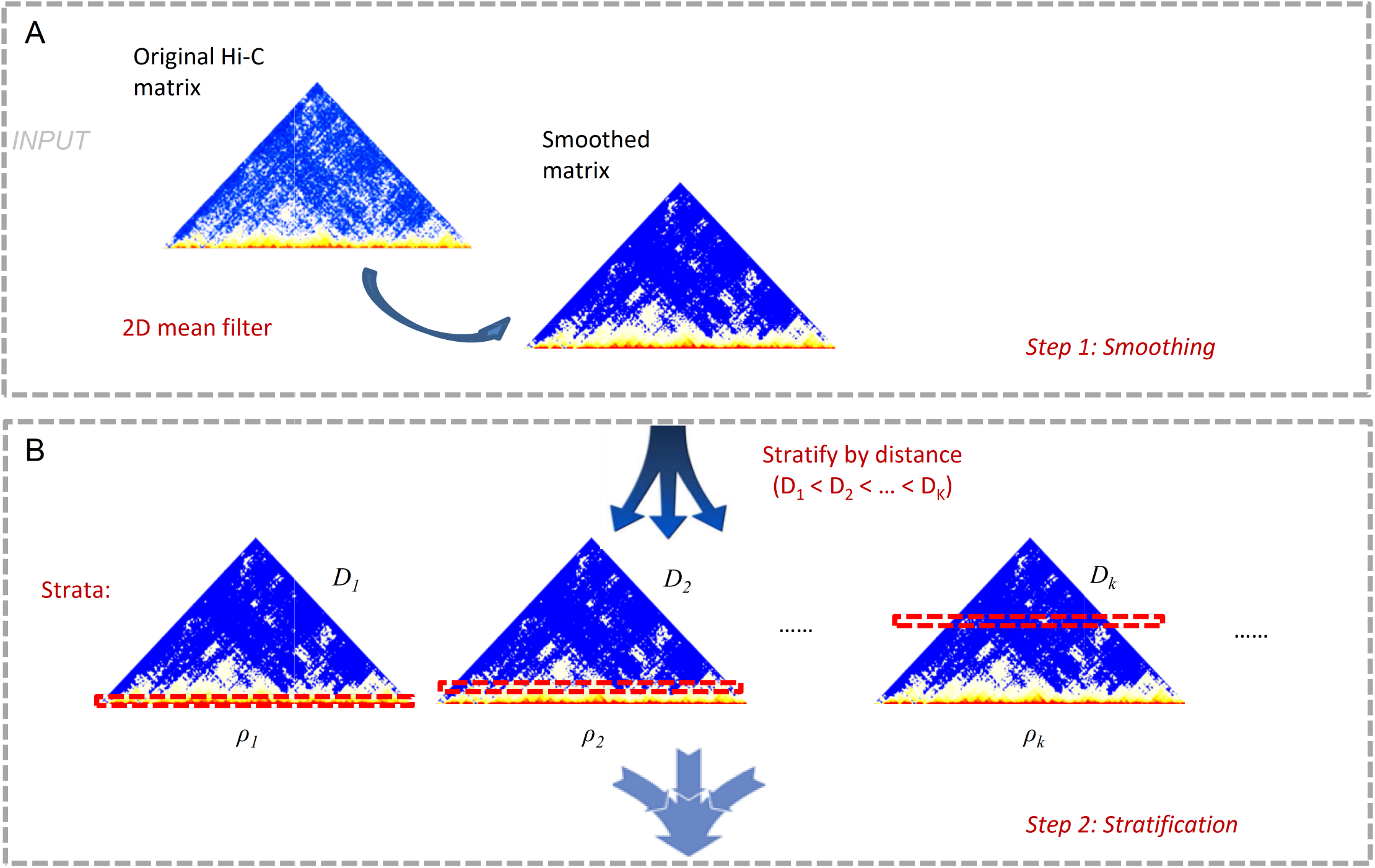
A schematic representation of our method.

### Distinguishing pseudo, real and non-replicates

We first evaluated the performance of our method on samples whose expected levels of reproducibility are known: pseudo-replicates (PR), biological replicates (BR) and non-replicates (NR). Biological replicates refer to two independent Hi-C experiments performed on the same cell types. Non-replicates refer to Hi-C experiments performed on different cell types. Pseudo replicates are generated by pooling reads from biological replicates together and randomly partitioning them into two equal portions. The difference between two pseudo-replicates only reflects sampling variation, without biological or technical variation. Therefore, we expect the reproducibility of pseudo-replicates to be the highest, followed by biological replicates and then non-replicates.

For testing, we first generated PR, BR and NR using Hi-C data in the hESC and IMR90 cell lines (Dixon et al. 2012) (details in Methods). We compared the performance of our method with Pearson correlation and Spearman correlation and investigated whether these metrics can distinguish PR, BR and NR (Fig. 3A and Supplemental Table S1). For the hESC dataset, our method correctly ranks the reproducibility of the three types of replicate pairs (PR>BR>NR), whereas Pearson and Spearman correlations both incorrectly rank BR lower than one or more of the NRs. For the IMR90 dataset, although all three methods infer the correct order of reproducibility, SCC separates BR from NR by a much larger margin (SCC: 0.19) than the other metrics (Pearson: 0.02 and Spearman: 0.03).

**Figure 3.**
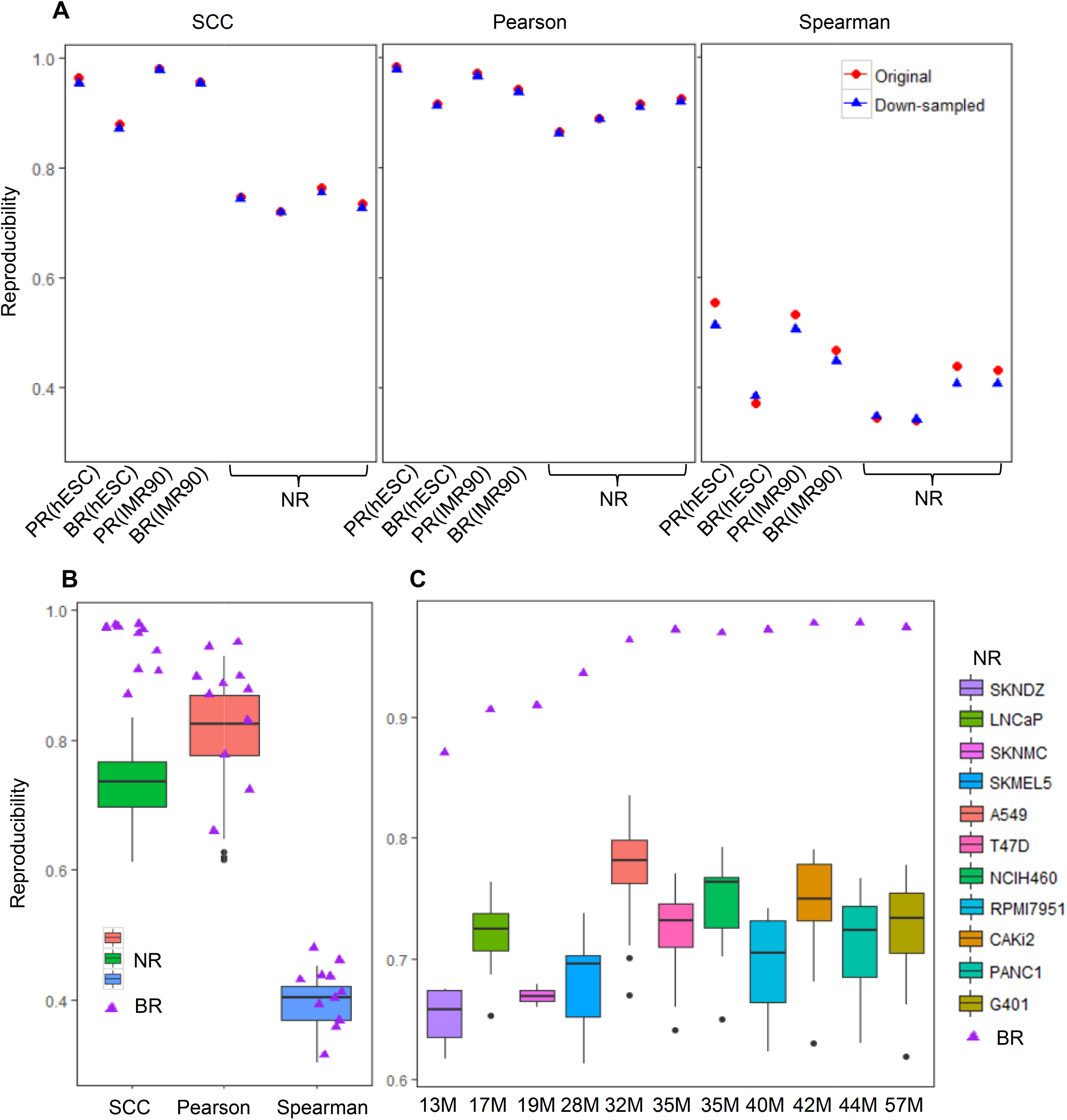
Discrimination of pseudo replicates (PR), biological replicates (BR) and non-replicates (NR). (A) Reproducibility scores for the illustration example (hESC and IMR90 cell lines) in Fig. 1. Red dots are the results in the original samples, and blue dots are the results after equalizing the sequencing depth in all samples. (B-C) Reproducibility scores for the BR and NR in the ENCODE 11 cancer cell lines. The triangle represents the score for a BR and the boxplot represents the distribution of the scores for NRs.(B) Reproducibility scores for BRs and NRs in all cell types. (C) SCC for BRs and the corresponding NRs in each cell type. From left to right, the cell lines are ordered according to the average sequencing depths of the biological replicates.

The sequencing depths differ substantially for the hESC (replicate 1: 60M; replicate 2: 271M) and IMR90 (replicate 1: 201M; replicate 2: 153M) datasets. To ensure that these differences were not confounding our evaluations, we subsampled all the replicates to 60 million reads and repeated the same analysis. As shown in Fig. 3A (blue dots), even with the same number of reads, Pearson and Spearman correlations still fail to distinguish real replicates from all non-replicates. On the contrary, our method consistently ordered the reproducibility of replicates correctly, indicating that it can capture the intrinsic differences between the samples, even those that differ in sequencing depth.

We expanded this analysis to a larger Hi-C dataset recently released by the ENCODE Project Consortium (The ENCODE Project Consortium, 2012). This dataset consists of Hi-C data from eleven cancer cell lines, with two biological replicates for each cell type (details are in Methods). For each cell type, we formed twenty non-replicate pairs with the remaining ten cell types and computed SCC, Pearson and Spearman correlations for BR and all NRs. As shown in Fig. 3B and Supplemental Table S2, SCC clearly distinguishes BRs from NRs (a p-value = 1.665 × 10^−15^, one-sided Kolmogorov-Smirnov test), while the other two methods fail to do so (Pearson: p-value = 0.084; Spearman: p-value = 0.254, K-S test). Because the sequencing depth of the Hi-C data varies across cell types, we also examined the separation between BRs and NRs for each cell type. As shown in Fig. 3C, SCC separates the BRs and NRs for all the cell types by a margin of at least 0.1, whereas the other two methods fail to separate them in more than half of the cell types (Supplemental Fig. S1). Furthermore, SCC illustrates a desirable correspondence to the sequencing depth. When the average sequencing depth between the biological replicates is relatively low (<30M), SCC monotonically increases with the sequencing depth; this behavior likely reflects insufficient coverage at the lower sequencing depths. In contrast, the value for SCC remains high and stable for greater sequencing depths (Fig. 3C), reflecting saturation of reproducibility and likely sufficient coverage. We investigate this property further in a later section.

### Evaluating biological relevance by constructing cell lineages

Next we used HiCRep to infer the interrelationship between cell types on a cell lineage. Because this assessment requires the reproducibility measure to quantify the subtle differences between closely related cells, it serves as a biologically relevant approach to evaluating the accuracy of the reproducibility measure. More importantly, it also evaluates the potential of our method as a measure for quantifying the similarities or differences of Hi-C matrices in different cell or tissue types.

For this analysis, we used the Hi-C data in human embryonic stem (ES) cells and in four cell lineages derived from them (Dixon et al. 2015), namely, mesendoderm (ME), mesenchymal stem cells (MS), neural progenitor cells (NP), and trophoblast-like cells (TB), with two biological replicates for each cell type. Using the A/B compartments in Hi-C data, Dixon et al. (2015) inferred the distance to the parental ES cell from the nearest to the farthest as ME, NP, TB and MS (Fig. 4A left). Importantly, the same relationships were also supported by our analysis of the previously published gene expression data (Xie et al. 2013) in the same cell types (Fig. 4A right).

**Figure 4.**
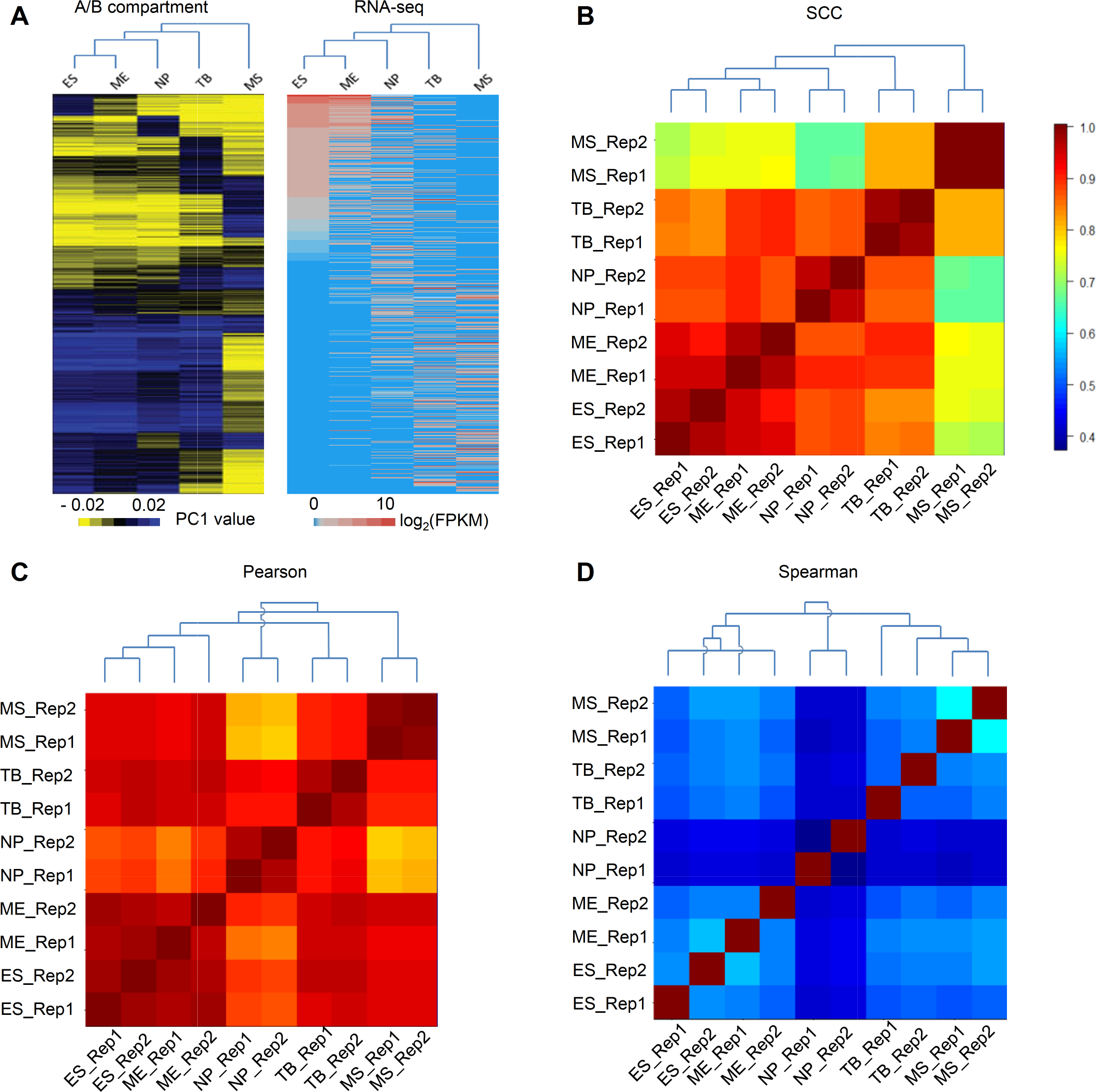
Estimating interrelationship between the ten samples in the human H1 ESC lineage. (A) The heatmap and lineage relationship between the ES cell and its five derived cells based on A/B compartments in Hi-C data (Dixon et al. 2015) and RNA-seq data in (Xie et al. 2013). (B-D) Estimated interrelationship based on the pairwise similarity score calculated using (B) SCC (C) Pearson correlation and (D) Spearman correlation. Heatmaps show the similarity scores. Dendrograms are resulted from a hierarchical clustering analysis based on the similarity scores. For easy visualization, the cell lines in the heatmaps are ordered according to their known distances to ES cells in (A). A decreasing trend of scores is expected from left to right (from bottom to top, respectively) if the estimated interrelationship agrees with the known lineage.

We first calculated the pairwise similarities between the ten samples (two replicates in each cell type) using SCC, Pearson and Spearman correlations (Supplemental Table S3). As shown in Supplemental Fig. S2, SCC again provided the best separation between real replicates and non-replicates among all three methods of comparison.

Next, we reconstructed the relationships among the cell lineages by performing hierarchical clustering based on the pairwise similarity scores. As shown in Fig. 4B, the dendrogram constructed based on SCC precisely depicts the interrelationships: all the biological replicates are grouped together as terminal clusters, and the relationships between cell lines exactly follow the tree structure in (Dixon et al. 2015) and (Xie et al. 2013) (Fig. 4A). The same results are obtained using the bias corrected interaction matrices (Supplemental Fig. 3A). In contrast, the dendrograms constructed based on Pearson (Fig. 4C) and Spearman correlation coefficients (Fig. 4D) group several non-replicates together and infer different relationships between some cell lines. For example, when using Pearson correlation, two ME replicates are not clustered together and NP is unexpectedly placed as the least related cell type to ES cells. When using Spearman correlation, an ES replicate is clustered with an ME replicate and again NP is unexpectedly predicted as the least related cell type to ES cells.

To further delineate how data smoothing contributes to HiCRep's performance, we also computed SCC on unsmoothed Hi-C matrices and Pearson and Spearman correlation coefficients on the smoothed Hi-C matrices obtained from our smoothing procedure. As shown in Supplemental Fig. S3B, we observed that the SCC analysis on unsmoothed data no longer recapitulates the expected relationships among cell lineages, indicating that the smoothing stage is an indispensable component of HiCRep. Furthermore, we observed that our smoothing procedure improves the performance of Pearson and Spearman based approaches (Supplemental Fig. S3 C-D), confirming its effectiveness. However, the improvement on Pearson and Spearman based approaches is not to the level achieved by SCC (Supplemental Fig. S3 C-D). For example, the tree based on Pearson incorrectly groups the ES cell and ME cell replicates together, and the tree based on Spearman incorrectly places the NP cell closest to ES cell. In addition, the Pearson correlation based on the smoothed matrices shows very little difference (range = (0.96, 1)) across cell lines, making it harder to distinguish closely related samples. Together, this indicates that neither SCC nor smoothing by itself can satisfactorily address the reproducibility issue, and both components are necessary for HiCRep to quantify biologically relevant differences between Hi-C contact maps.

We further expanded this analysis using the recently published Hi-C data in fourteen human primary tissues and two cell lines (Schmitt et al. 2016) (Supplemental Table S4). Because biological replicates are not available for all the samples, our analysis focused on quantifying the relationships between tissues or cells. Again, the lineage constructed based on SCC reasonably depicted the tissue and germ layer origins of the samples (Fig. 5A): hippocampus and cortex were grouped together; right ventricle and left ventricle were grouped together; endodermal tissues such as pancreas, lung, and small bowel were placed in the same lineage. Neither Pearson nor Spearman correlation performed as well as SCC did. For example, right and left ventricles were not grouped together by Spearman correlation (Fig. 5C). These results confirm the potential of our method as a measure for quantifying the difference in Hi-C data between cell or tissue types.

**Figure 5.**
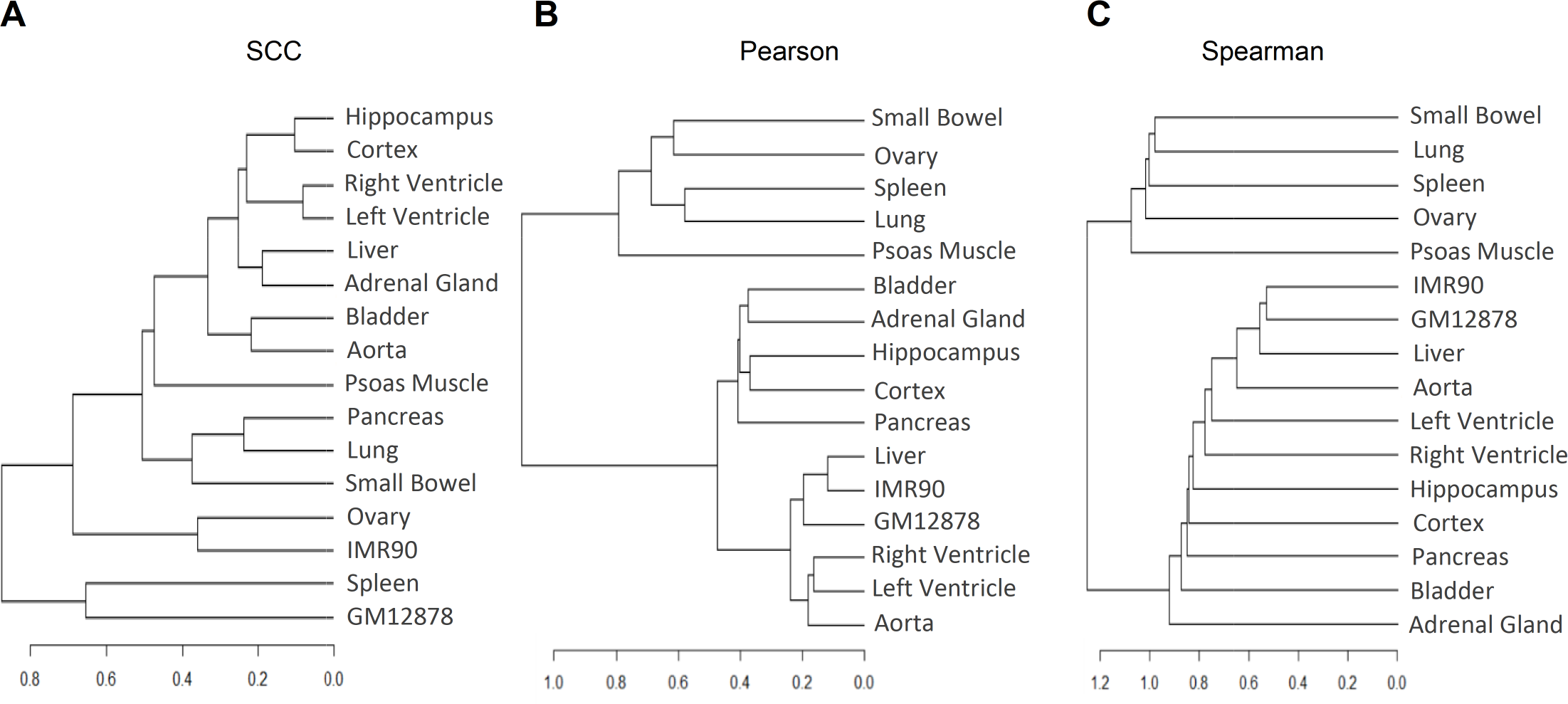
Estimated interrelationship for fourteen human primary tissues and two cell lines in (Schmitt et al. 2016). The dendrograms are resulted from a hierarchical clustering analysis based on the pairwise similarity calculated using (A) SCC, (B) Pearson correlation and (C) Spearman correlation.

### HiCRep is robust to different choices of resolution

Depending on the sequencing depth, Hi-C data analysis may be performed at different resolutions. A good reproducibility measure should perform well despite the choice of resolution. To evaluate the robustness of our method, we repeated the clustering analysis for the human ES and ES-derived cell lineages using data processed at several different resolutions (i.e., 10Kb, 25Kb, 40Kb, 100Kb, 500Kb, 1Mb). Again, as shown in Fig. 6 and Supplemental Table S5, SCC accurately inferred the expected relationships between ES and its derived cell lines at all resolutions considered, whereas Pearson and Spearman correlations inferred the expected relationships only at 500Kb and 1Mb. Furthermore, unlike Pearson and Spearman correlations, whose values drastically change at different resolutions, the values of SCC remain in a consistent range across all resolutions. The complete trees inferred by SCC at different resolutions (Supplemental Fig. S4) all agree with the expected relationship. These results confirm the robustness of our method to the choice of resolution.

**Figure 6.**
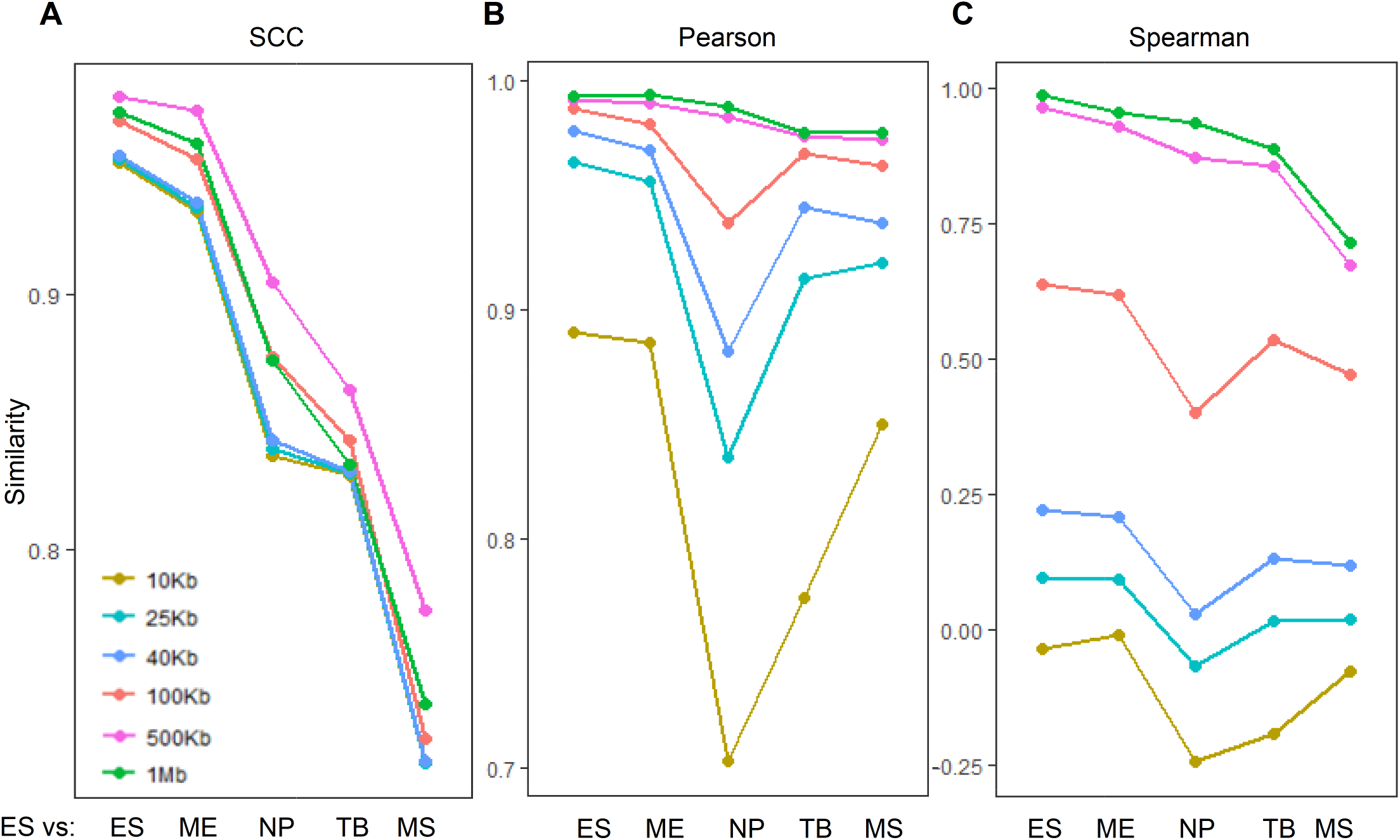
Estimated similarity between the human H1 ES cell and its derived cells at different resolutions. (A) SCC, (B) Pearson correlation coefficient, and (C) Spearman correlation coefficient.

### Detecting differences in reproducibility due to sequencing depth

Sequencing depth is known to affect the signal-to-noise ratio and the reproducibility of Hi-C data (Lajoie et al. 2015). Insufficient coverage can reduce the reproducibility of a Hi-C experiment. As a quality control tool, a reproducibility measure is expected to be able to detect the differences in reproducibility due to sequencing depth. To evaluate the sensitivity of our method to sequencing depth, we downsampled all the samples in the H1 ES cell lineage (Dixon et al. 2015) to create a series of subsamples with different sequencing depths (25%, 50% and 75% of the original sequencing depth). We then computed SCC for all subsamples. As shown in Fig. 7A and Supplemental Table S6, SCC monotonically decreases with sequencing depth in all data sets. This confirms that our method can reflect the change of reproducibility between replicate experiments due to sequencing depth.

**Figure 7.**
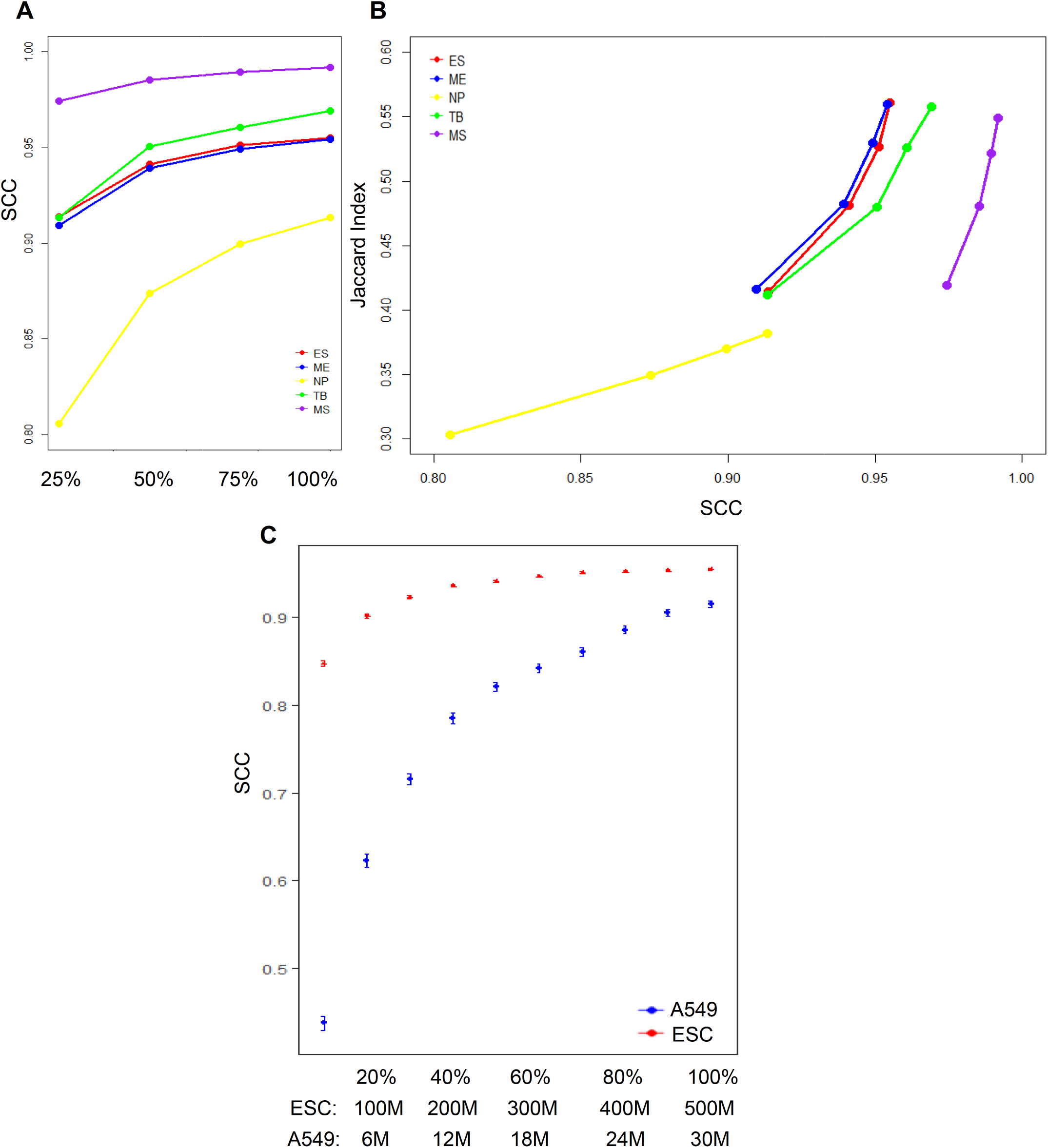
Detecting the change of reproducibility due to sequencing depth using SCC. (A) SCC of downsampled biological replicates (25%, 50%, 75%, 100% of the original sequencing depth) for the five cell lines on the H1 ES cell lineage. (B) Relationship between SCC and Jaccard index, which measures the proportion of shared significant contacts identified by Fit-Hi-C between replicates for samples in (A).(C)Saturation curves of SCC for datasets with different coverages. Plotted is the SCC at different subsamples (10%-90%) of the original samples with 90% confidence intervals. The blue dots represent H1 human ESC data (original sequencing depth=500M). The red dots represent the A549 data (original sequencing depth=30M).

Furthermore, we investigate whether the reproducibility between Hi-C experiments inferred by SCC reflects the reproducibility at the level of significant contacts. To proceed, we identified significant contacts in these subsamples using Fit-Hi-C (q-value cutoff=0.05) (Ay et al. 2014). For each subsample, we computed the reproducibility at the level of significant contacts using the Jaccard index, i.e. the ratio of the number of shared significant contacts over the number of significant contacts identified in either replicate. As shown in Fig. 7B and Supplemental Table S6, the Jaccard index monotonically increases with the SCC score in each cell line. In addition, the cell line (NP), which has a significantly lower SCC score than other cell lines also shows a significantly lower level of shared significant contacts. This confirms that our method can also reflect the change of reproducibility due to sequencing depth at the level of significant contacts.

### Guiding the selection of the optimal sequencing depth

Having established that SCC can reflect the change of reproducibility due to the change of sequencing depth, we propose to use the saturation of SCC as a criterion to determine the most cost-effective sequencing depth that achieves a reasonable reproducibility. To illustrate how to use our method to determine the optimal sequencing depth, we created subsamples at a series of reduced sequencing depths from the Hi-C data in the H1 ES cell in (Dixon et al. 2015) (original depth=500M) by down-sampling. As shown in Fig. 7C and Supplemental Table S7, SCC initially increases rapidly with the increase of sequencing depth when the number of total reads is less than 200 million (slope of line from 10% to 40% depth is 0.0591 per 100M). It continues to increase at a reduced rate (slope of line from 40% to 70% depth is 0.01 per 100M), and eventually reaches a plateau (slope of line from 70% to 100% depth is 0.002 per 100M). To determine the lowest sequencing level that achieves similar reproducibility as the original data, we computed the 99% confidence intervals of SCC at each sequencing depths. Starting at 350M (70% of the original depth), the confidence intervals of SCC between two adjacent depth levels overlap with each other and the difference of SCC from that of the original depth is less than 0.004. This indicates that the reduced samples can achieve a similar level of reproducibility as the original one by using about 70% of the original depth for this dataset. Further increase of sequencing depth beyond this point does not significantly improve reproducibility.

As a comparison, we performed a similar analysis using a dataset with relatively low sequencing depth (30M Hi-C reads from the A549 cell line). We observe that all the reduced samples have a significantly lower reproducibility than the original sample at the 99% significance level and show a steep increase of SCC throughout all downsample levels (Fig. 7C and Supplemental Table S7). From the 90% depth to the original depth, there is still an increase of SCC of 0.01, compared with 0.0015 for the hESC dataset, suggesting that this dataset may not reach saturation in reproducibility at its original sequencing depth. For this dataset, further increase of sequencing depth may improve reproducibility.

## Discussion

Although there has been a dramatic increase in the scope and complexity of Hi-C experiments, analytical tools for data quality control have been lacking. Current approaches for assessing Hi-C data reproducibility may lead to incorrect conclusions because they fail to take into consideration the unique spatial characteristics of Hi-C data. In this work, we developed a new method, HiCRep, for assessing the reproducibility of Hi-C contact frequency maps. By effectively taking account of the spatial features of Hi-C data, our reproducibility measure overcomes the limitations of Pearson and Spearman correlations and can differentiate the reproducibility of samples at a fine level. The empirical evaluation showed that our method distinguished subtle differences between closely related cell lines, biological replicates and pseudo replicates, and it produced robust results at different resolutions.

The SCC statistic has several properties that make it well-suited as a reproducibility measure for providing standardized, interpretable, automatable and scalable quality control. First, this statistic has a fixed scale of [−1, 1], which makes it easy to standardize the quality control process and compare reproducibility across samples. Second, it is intuitive and easy to interpret. It can be interpreted as a weighted average correlation coefficient over different interaction distances. This straightforward interpretation makes it accessible to experimentalists. Third, our statistic is fast to compute and is directly applicable to the raw contact matrix. It is easily scalable for monitoring data quality for a large number of experiments. Furthermore, we also provide an estimator for the variance of this statistics, such that the statistical significance of the difference in reproducibility can be inferred. Using this estimator, we establish a procedure to determine the sufficiency of sequencing depth.

In summary, we develop a novel method to accurately evaluate the reproducibility of Hi-C experiments. The presented method is a first step toward ensuring high reproducibility of Hi-C data. We also show that this method can be used as a similarity measure for quantifying the differences in Hi-C data between different cell and tissue types. Thus, HiCRep is a valuable tool for the study of 3D genome organization.

## Methods

### Datasets

The datasets analyzed in this study were obtained from the public domain, as described below. Hi-C data sets used in this project can be visualized in the 3D genome browser (http://3dgenome.org).

We obtained the Hi-C data of human embryonic stem cells (hESCs) and human IMR90 fibroblasts from (Dixon et al. 2012) (GEO accession number: GSE35156). Each cell type has two biological replicates.

We obtained the Hi-C data of human embryonic stem (ES) cells and four human ES-cell-derived lineages, mesendoderm (ME), mesenchymal stem (MS) cells, neural progenitor (NP) cells and trophoblast-like (TB) cells from (Dixon et al. 2015) (GEO accession number: GSE52457). Each cell type has two biological replicates.

We obtained the Hi-C data of eleven human cancer cell lines from the ENCODE data portal (Sloan et al. 2016) (https://www.encodeproject.org/matrix/?type=Experiment&status=released&assay_slims=3D+chromatin+structure&award.project=ENCODE&assay_title=Hi-C). This dataset was produced by Dekker lab. It includes cell lines of G401, A549, CAKi2, PANC1, RPMI7951, T47D, NCIH460, SKMEL5, LNCaP, SKNMC and SKNDZ. Each cell line has two biological replicates. The sequencing depths of the datasets can be found in Supplemental Table S8.

We obtained the Hi-C data of fourteen human primary tissues from (Schmitt et al. 2016) and (Leung et al. 2015). The tissues include adrenal gland (GSM2322539), bladder (GSM2322540, GSM2322541), dorsolateral prefrontal cortex (GSM2322542), hippocampus (GSM2322543), lung (GSM2322544), ovary (GSM2322546), pancreas (GSM2322547), psoas muscle (GSM2322551), right ventricle (GSM2322554), small bowel (GSM2322555), spleen (GSM2322556), liver (GSM1419084), left ventricle (GSM1419085), and aorta (GSM1419086). The tissues were collected from four donors, each of which provides a subset of tissues. To minimize variation due to individual difference, we used the samples from the two donors with the largest number of tissues. If one tissue sample consists of multiple replicates from a single donor, the replicates were merged into a single dataset. We obtained the GM12878 cell data from (Selvaraj et al. 2013) (GSM1181867, GSM1181867) and the IMR90 cell data from (Dixon et al. 2012) (GSM862724, GSM892307).

### Data Preprocessing

We generated the Hi-C contact matrix using the pipeline from (Dixon et al. 2015). Briefly, the paired-end reads were first aligned to the hg19 reference genome assembly using BWA (Li and Durbin 2009). The unmapped reads were filtered, and potential PCR duplicates were removed using Picardtools (https://broadinstitute.github.io/picard/). We analyzed Hi-C reads mapped to human genome assembly hg19 because many of the published datasets were mapped to this assembly and this assembly has the deepest annotation of candidate functional noncoding sequences. While the human genome assembly GRCh38 has improved contiguity in local regions, both assemblies are high quality. Thus genome-wide analyses such as our assessments of reproducibility are not expected to differ significantly when the Hi-C reads are aligned to different assemblies. Importantly, our evaluation of reproducibility is not dependent on the exact locations to which reads map, but rather it uses the distances between two interacting sequences. Thus our metrics should be stable across similar assemblies.

For most analysis, we used 40kb bins. To obtain contact maps at this resolution, we divided the genome into 40kb bins as in (Dixon et al. 2015) and obtained the interaction frequency by counting the number of reads falling into each pair of bins.

Our analysis only considered the intra-chromosomal interactions and only used the contacts within the range of 0-5Mb in the reproducibility assessment. This range was chosen based on the observation that most of the A/B compartments have an interaction size of about 5Mb, and interactions over 5Mb in distance are rare (< 5% of reads) and highly stochastic. To evaluate the effect of this parameter, we constructed the ES and its derived cell lineages using interactions in several ranges, including 0-4Mb, 0-5Mb, 0-6Mb, 0-8Mb, and 0-10Mb (Fig. 4B for 0-5Mb and Supplemental Fig. S5 for others). The ranges of 0-4Mb, 0-5Mb and 0-6Mb gave the best results, confirming that 0-5Mb is a reasonable choice. Only the bins with at least one count in at least one of the samples are kept for computing Pearson and Spearman correlations. All the datasets were preprocessed using the same procedure.

Our method is applicable to both raw and bias corrected data. However, to ensure the reproducibility assessment is free of assumptions made in the bias correction procedures (Imakaev et al. 2012; Hu et al. 2012; Cournac et al. 2012) and faithfully reflects the nature of the raw data, we chose to apply our method directly to raw data without bias correction. For the ES and its derived cell lines, we also applied our method to the biased corrected matrices, in addition to the raw data, as a comparison. The bias correction was performed using the Iterative Correction and Eigenvector decomposition procedure (ICE) (Imakaev et al. 2012).

### 2D mean filter smoothing

Because the space of interactions surveyed by Hi-C experiments is very large, achieving sufficient coverage is still challenging. When samples are not sufficiently sequenced, the local variation introduced by under-sampling can make it difficult to capture large domain structures.

To handle this issue, we first smooth the contact map before assessing reproducibility. Although smoothing will reduce the individual spatial resolution, it can improve the contiguity of the regions with elevated interaction, consequently enhancing the domain structures. It has been found effective in commonly-used Hi-C normalization methods (Imakaev et al. 2012; Yaffe and Tanay 2011).

We use a 2D mean filter to smooth the contact map. The filter replaces the read count of each contact in the contact map with the mean counts of all contacts in its genomic neighborhood. This filter is fast to compute and is effective for smoothing rectangular shapes (Davies 2012) like domain structures in Hi-C data. Specifically, let *C_n×n_* denote a *n*×*n* contact map and *c_ij_* denote the counts of the interaction between loci *i* and *j*. Given a span size *h*>0, the smoothed contact map after passing an *h^th^* 2D mean filter is defined as follows:

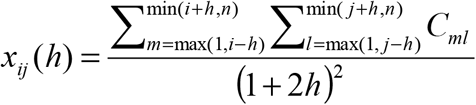

A visualization of the smoothing effect with different window sizes is shown in Supplemental Fig. S6,

### Selection of smoothing parameter

The span size *h* is a tuning parameter controlling the smoothing level. A very small *h* might not reduce enough local variation to enhance the boundaries of domain structures, while a large *h* will make the boundaries of domain structures blurry and limit the spatial resolution. Therefore, the optimal *h* should be adaptively chosen from the data.

To select *h* objectively, we developed a heuristic procedure to search for the optimal choice. Our procedure is designed based on the observation that the correlation between contact maps of replicate samples first increases with the level of smoothness and then plateaus when sufficient smoothness is reached. To proceed, we used a pair of reasonably deeply sequenced interaction maps as the training data. We randomly sampled 10% of the data ten times. For each subsample, we computed the stratum-adjusted correlation coefficient (SCC, described in a later section) at a series of *h*'s in the ascending order and recorded the smallest *h* at which the increment of SCC was less than 0.01. The mode of *h* among the ten subsamples was selected as the final span size. The detailed steps are shown in Algorithm 1 (Supplemental File S1).

Because the level of local variation in a contact map depends on the resolution used to process the data, the span size required to achieve sufficient smoothness varies according to resolution. Hence, a proper *h* for each resolution needs to be trained separately. However, at a given resolution, it is desirable to use the same *h* for all datasets, so that the downstream reproducibility assessment can be compared on the same basis. To reduce the chance of over-smoothing due to sparseness caused by insufficient coverage when training *h*, we used a deeply sequenced data set as training data.

Here we obtained *h* in our analysis from the Human H1 ESC dataset (Dixon et al. 2015). This dataset was deeply sequenced (330M and 740M reads for its two replicates) and had a reasonable quality (Dixon et al. 2015), making it suitable as training data. We processed the data using a series of resolutions (10Kb, 25Kb, 40Kb, 100Kb, 500Kb and 1Mb), and then selected *h* for each resolution using the procedure described above. We obtained *h*=20, 11, 5, 3, 1, and 0 for the resolution of 10Kb, 25Kb, 40Kb, 100Kb, 500Kb and 1Mb, respectively. These values were used throughout our study for all datasets at the corresponding resolutions. The robustness of our procedure was assessed using the Human H1 ESC dataset and four derived cell lines (details are in the Results section).

### Stratification by distance

To take proper account of the distance effect in reproducibility assessment, we stratify the contacts by the genomic distance between their interaction loci. Specifically, let *X_n×n_* be an *n×n* smoothed contact map at a resolution of bin size *b*. We compute the interaction distance for each contact *x_ij_* as *d_ij_* = |*j - i|×b* and then stratify the contacts by *d_ij_* into *K* strata, *X_k_* = {*x_ij_: (k-1)b < d_ij_ ≤ kb}*, *k = 1*, *… K*. Here we consider the interaction distance of 0-5Mb. This leads to *K* = 125 for the bin size *b* = 40kb. If *x_ij_* is 0 in both samples, then it is excluded from the reproducibility assessment.

### Stratum-adjusted correlation coefficient (SCC)

Our reproducibility statistic is motivated from the generalized Cochran-Mantel-Haenszel (CMH) statistic M^2^. The CMH statistic is a stratum-adjusted summary statistic for testing if two variables are associated while being stratified by the third variable (Agresti 2012), for example, the association between treatment and response stratified by age. Though originally developed for categorical data, it is also applicable to continuous data (Mantel 1963) and can detect consistent linear association across strata. However, the magnitude of M^2^ depends on the sample size; therefore, it cannot be used directly as a measure of the strength of the association. When there is no stratification, the CMH statistic is related to the Pearson correlation coefficient *ρ* as M^2^ = *ρ^2^* (*N*−1), where *N* is the number of observations (Agresti 2012). This relationship allows the strength of association summarized by M^2^ to be represented using a measure that has a fixed scale and is comparable across different samples. However, *ρ* does not involve stratification. This motivates us to derive a stratum-adjusted correlation coefficient (SCC) to summarize the strength of association from the CMH statistic when there is stratification.

### Derivation of stratum-adjusted correlation coefficient (SCC)

Let (*X*, *Y)* denote a pair of samples with *N* observations. The observations are stratified into *K* strata, and each stratum has *Nk* observations such that 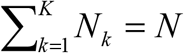 Denote the observations in stratum *k* as (*X_1_k__*, *Y_1_k__*), …., (*X_N_k__, Y_N_k__*) and the corresponding random variables as (*X_k_, Y_k_*) respectively. In our context, (*X_i_k__, Y_N_k__*) are the smoothed counts of the *i*^th^ contact on the *k*^th^ stratum in the two contact maps *X* and 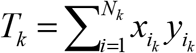 the CMH statistics is defined as

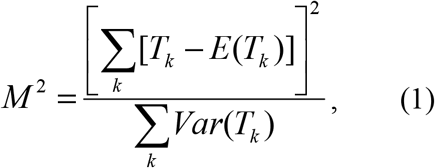

Where *E*(*T_k_*) and *Var* (*T_k_*) are the mean and variance of *T_k_* under the hypothesis that *X_k_* and *Y_k_* are conditionally independent given the stratum,

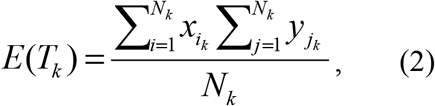

and

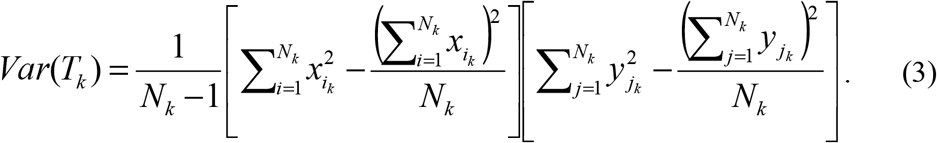

To derive the stratum-adjusted correlation coefficient from the CMH statistic, write the Pearson correlation coefficient *ρ_k_* for the *k*th stratum as 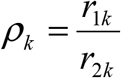, where

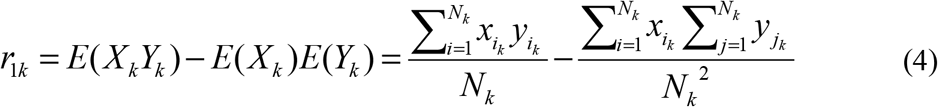

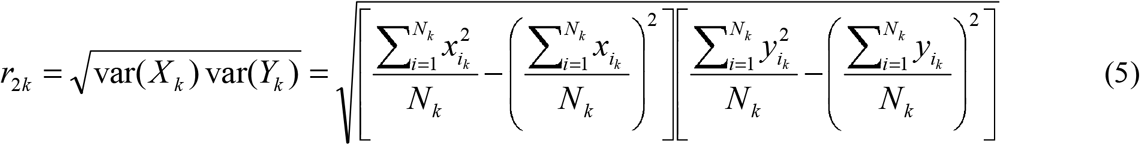

It is easy to see that 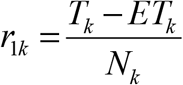 and 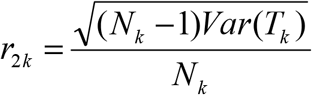. Then we can represent M^2^ using *ρ_k_*,

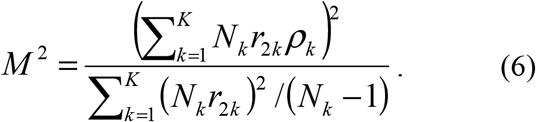

Define

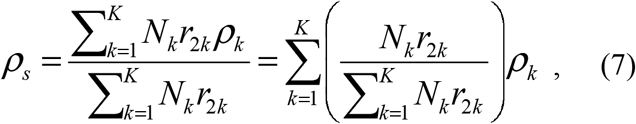

then

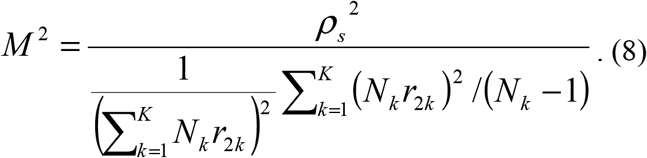

From (8), it can be seen that 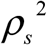 reflects the strength of association in M^2^. This strength relates to M^2^ in a similar way as the Pearson correlation to M^2^ in the case without stratification. As shown in (7) *ρ_s_* is a weighted average of the stratum-specific correlation coefficients, with weights 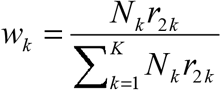 assigned according to the variance and sample size of a stratum. We call ***ρ_s_*** the stratum-adjusted correlation coefficient (SCC). Similar to standard correlation coefficients, it satisfies −1≤ *ρ_s_* ≤1. A value of *ρ_s_* = 1 corresponds to a perfect positive correlation, a value of *ρ _s_* = 1 corresponds to a perfect negative correlation, and a value of *ρ_s_* = 0 corresponds to no correlation.

The variance of *ρ_s_* can be computed as

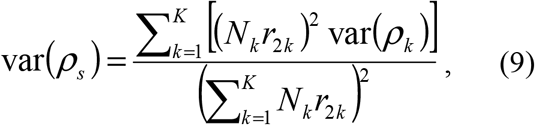

where (*ρ_k_*) is the asymptotic variance for the Pearson correlation coefficient in a single stratum and can be computed using Fisher transformation (Fisher 1921) as follows. Let *r_k_* be the sample version of *ρ_k_* and *Z_k_* be the Fisher transformation of 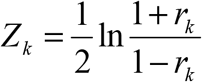 then *Z_k_* is approximately normal distributed with mean 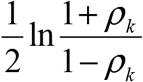 and variance 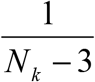 (Fisher 1921). By the Delta method (Casella and Berger 1993), *r_k_* is asymptotically normally distributed with mean *ρ^k^* and the asymptotic variance as 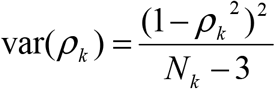.

The idea of obtaining an average correlation coefficient based on the CMH statistic has been explored in (Rubenstein and Davis 1999) in the context of contingency tables with ordered categories. However, its derivation has several errors, which lead to a different statistic that ignores the sample size differences in different strata.

### Variance stabilized weights

The downside for Equation (7) is that it is based on the implicit assumption in the CMH statistic that the dynamic ranges of X and Y are constant across strata. However, in Hi-C data, the read counts for contacts with short interaction distances have a much larger dynamic range than those with long interaction distances. As a result, the weights for the strata with large dynamic ranges will dominate (7), due to the large values of their *r_2k_*. To normalize the dynamic range, we rank the contact counts in each stratum separately and then normalize the ranks by the total number of observations *N_k_* in each stratum, such that all strata share a similar dynamic range. We then *r_2k_* compute in the weights in (7) and (9) using the normalized ranks, instead of the actual counts, i.e.

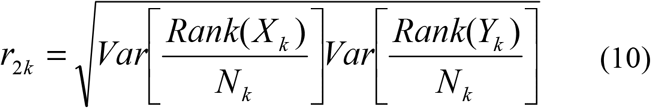

The stratum-specific correlation *ρ_k_* still computed using actual values rather than ranks, as actual alues have better sensitivity than ranks when there are a large number of low counts.

### Software availability

We have implemented our method as an R package (R Core Team 2016). It is publicly available as the HiCRep package on GitHub (https://github.com/qunhualilab/hicrep) and the Supplemental Materials.

## Acknowledgements

The authors thank Dekker's lab for providing the unpublished cancer cell line Hi-C data for our analysis. This work was supported by NIH training grant T32 GM102057 (CBIOS training program to The Pennsylvania State University), a Huck Graduate Research Innovation Grant to TY, and by NIH grants R01GM109453 (to QL), U01CA200060 (to FY), R24DK106766 (to RCH and FY), U54HG006998 (to RCH), and U41HG007000 (to WSN).

## Authors' contributions

QL and FY designed the study. QL and TY formulated the problem and developed the computational method and the SCC statistic. FZ provided assistance in the formulation of SCC statistic. QL, FY and TY designed the data analysis and performance evaluation. GY and WN designed the data analysis for ENCODE cancer cell data. TY implemented the method, developed the software package and analyzed the data. TY and FS optimized the software package. QL, TY, and FY interpreted the results. RH provided assistance on the biological interpretation of the results. TY, RH, FY and QL wrote the paper. All authors read and approved the final manuscript.

